# Consistent EMG encoding during reach and grasp by neurons in motor cortex

**DOI:** 10.64898/2026.03.02.709159

**Authors:** Chiara Ciucci, Xuan Ma, Fabio Rizzoglio, Eric Gasper, Anton R. Sobinov, Lee E. Miller

**Author notes:** Indicates equal contributions (Chiara Ciucci, Xuan Ma).

## Abstract

Identifying what the activity of neurons in the primary motor cortex (M1) represents is essential for understanding motor computation. Yet the relation of M1 to muscles and movement remains unresolved, in part because the kinematic variables M1 appears to encode differ across the limb: hand velocity during reaching, but joint position during grasping. This discrepancy has been taken to imply fundamentally distinct control strategies for proximal and distal segments. Yet, here we show that a single, muscle-based control principle accounts well for both observations. We recorded neural activity, electromyographic (EMG) signals, and movement kinematics from macaque monkeys performing planar reaching, wrist movement, or free-form grasping tasks. Across behaviors, EMG-based encoding models explained M1 firing rates as accurately as the best-performing kinematic model: velocity for reaching, position for grasping, and either variable for wrist movements. Impulse response analysis revealed that these task-dependent kinematic relationships arise from the differences in mechanical impedance of each limb segment: the inertial and intersegmental coupling which dominate the dynamics of the proximal arm are far less important in the hand. These findings indicate that M1 output has a relatively simple, consistent relation to muscle activation and that the apparent divergence in kinematic encoding is a consequence of limb mechanics rather than distinct cortical strategies.

## Introduction

Understanding the language of the neural activity in the motor cortex (M1) is essential for interpreting the computations that lead to arm movement, from reaching to dexterous object manipulation. The neuroscientific community has been attempting to decipher this language for decades, beginning from the first recordings of single neurons from awake monkeys in the mid-1960s ^1^. In those early experiments, Evarts found a group of neurons which were more closely related to applied force or muscle activity than they were to the kinematics of wrist movements. However, those neurons were a small minority, compared to the much larger group of neurons he could analyze only qualitatively, which were driven more strongly when the monkey grasped the device than by subsequent wrist movements. In later experiments which presaged the modern brain-computer interface (BCI), Humphrey was able to decode multiple wrist movement related variables from small numbers of M1 neurons; force most accurately, but angular velocity nearly as well, both of which surpassed the accuracy of position decoding ^2^. The classic reaching studies of Georgopoulos ^3^, found motor cortex discharge to be modulated by the direction of the movement, rather than the final position of the hand. That basic observation was reinforced by many subsequent experiments studying reaching movements ^4-7^ and laid the foundation for velocity-based decoding that has now become nearly ubiquitous in modern BCIs.

However, recent quantitative studies of grasping have reported a much stronger correspondence between M1 neurons and joint angles than with angular velocity, revealing an apparent discord between the representation of the proximal and distal limb ^8-10^. A possible explanation is that arm and hand movements are represented fundamentally differently at the level of M1, perhaps relying on different processing within the spinal cord ^11-13^. We hypothesized, instead, that neurons in M1 speak the language of muscle activity; the conflicting neural dynamics across tasks simply reflect the fundamental physical difference between the proximal arm, dominated by inertial and velocity-related coupling forces ^14^, and the hand, for which elastic forces and the length-tension properties of muscle ^15,16^ may play a greater role.

In this study, we directly compare the motor cortical encoding of kinematic – position and velocity – and EMG variables as monkeys performed reaching, wrist movement, or grasping tasks. We found differing optimal lags between each of these classes of variables and neural firing rates and replicated the fundamental observations of the classic grasping and reaching studies as well as the more recent kinematic grasp encoding studies. Ultimately, we looked for differences in mechanical limb impedance that might unify muscle-encoding across the arm and hand.

## Results

We recorded single-neuron activity from six monkeys using multi-electrode arrays implanted in either the hand (Monkeys PO, TO, GR) or arm area (Monkeys AR, CH, and MH) of M1 (See Extended Data Table 1). We also collected EMGs from arm and hand muscles of all monkeys except CH and MH (Extended Data Table 2). The data from CH and MH are used to support the main conclusions drawn from AR. We recorded the data in three types of tasks: object grasp, wrist movement, and reach. PO and TO performed the grasp task with the arm comfortably positioned in front of the body and the upper arm and forearm restrained (Figure 1A). During the task, the experimenter repeatedly presented one of several objects in various positions and orientations to promote diverse grasp configurations and reduce inter-joint correlations during repeated grasp- and-release movement. The monkeys’ movements were tracked using four-camera markerless motion tracking.

**Figure 1:**
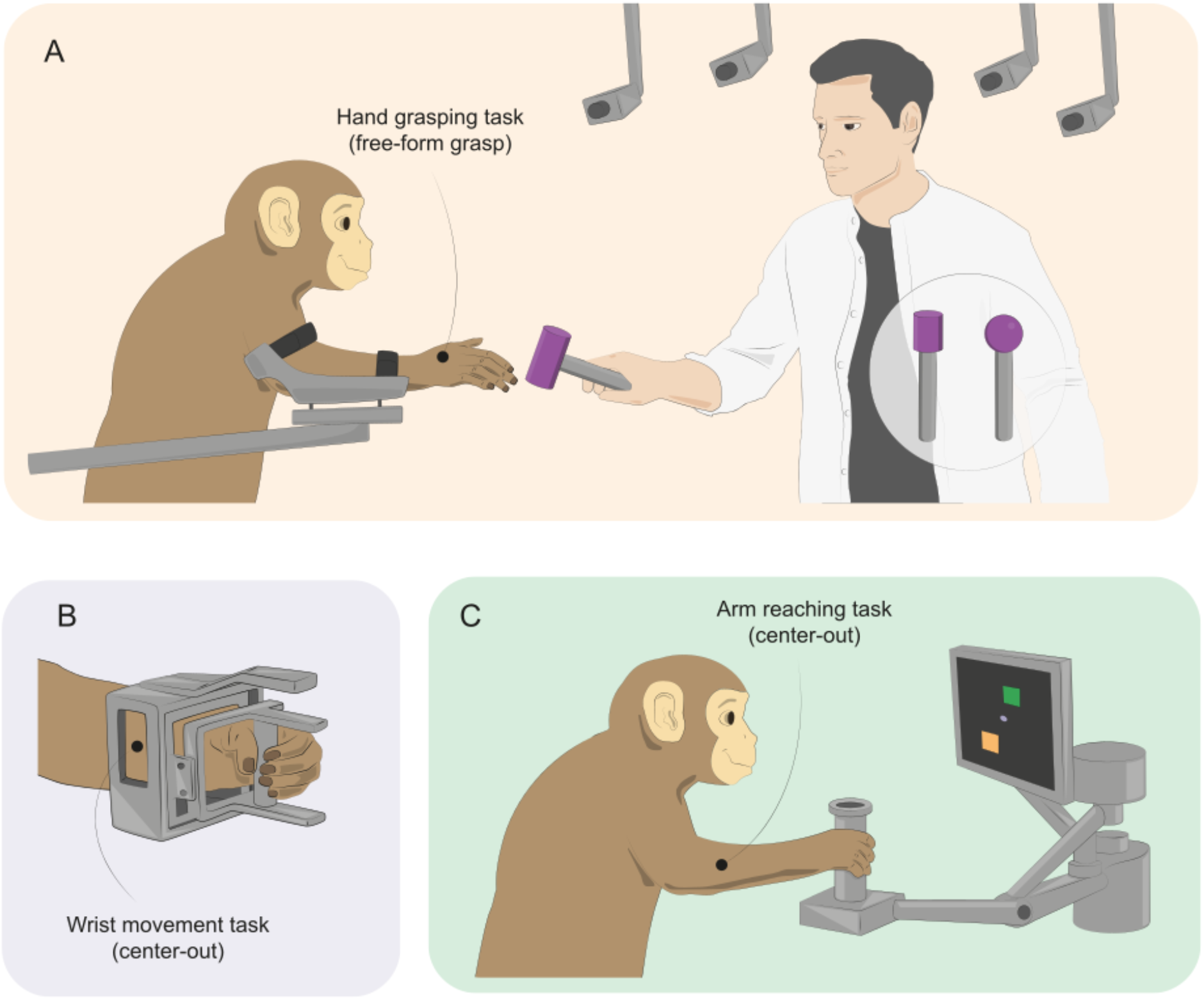
Schematic representations of the three behavioural tasks. (A) Grasp task, with the monkey’s upper arm and forearm restrained, allowing only wrist and finger movements. The experimenter presented objects of different shapes near the hand to elicit diverse grasp-and-release movements. (B) Wrist movement task, in which the monkey grasped a wrist-operated manipulandum with the forearm restrained and made flexion–extension and radial–ulnar deviation movements to move a cursor from a central target to one of eight outer targets. (C) Reach task -- the monkey grasped the handle of a planar manipulandum with the upper arm largely in the sagittal plane and made reaching movements in the horizontal plane to eight outer targets. Movement kinematics were captured either with a multi-camera markerless motion tracking system (Grasp) or encoders mounted on the manipulandum (Wrist and Reach).

In addition to the grasp task, we collected data from monkey GR during a wrist center-out task using the device pictured in 1B. The monkey controlled the movement of a cursor on a screen from a central target to one of eight randomly selected outer targets arranged in a circle around the center, while we monitored flexion/extension and radial/ulnar deviation movements of the wrist. Finally, we recorded data from monkeys AR, CH, and MH during a planar center-out reach task. The monkey grasped the handle of a planar manipulandum (Figure 1C) and made movements in the horizontal plane to move a cursor as in the wrist task.

Figure 2 shows a representative segment of data collected from monkey TO during the grasp task. Raster diagrams at the top show the discharge of the 10 neurons that responded most strongly during this session. Immediately below are EMG signals from four of the 16 muscles recorded from this monkey. At the bottom of the figure are three of the 24 kinematic degrees of freedom we extracted from the motion capture data. We collected analogous data from the other monkeys during the reach and wrist tasks.

**Figure 2:**
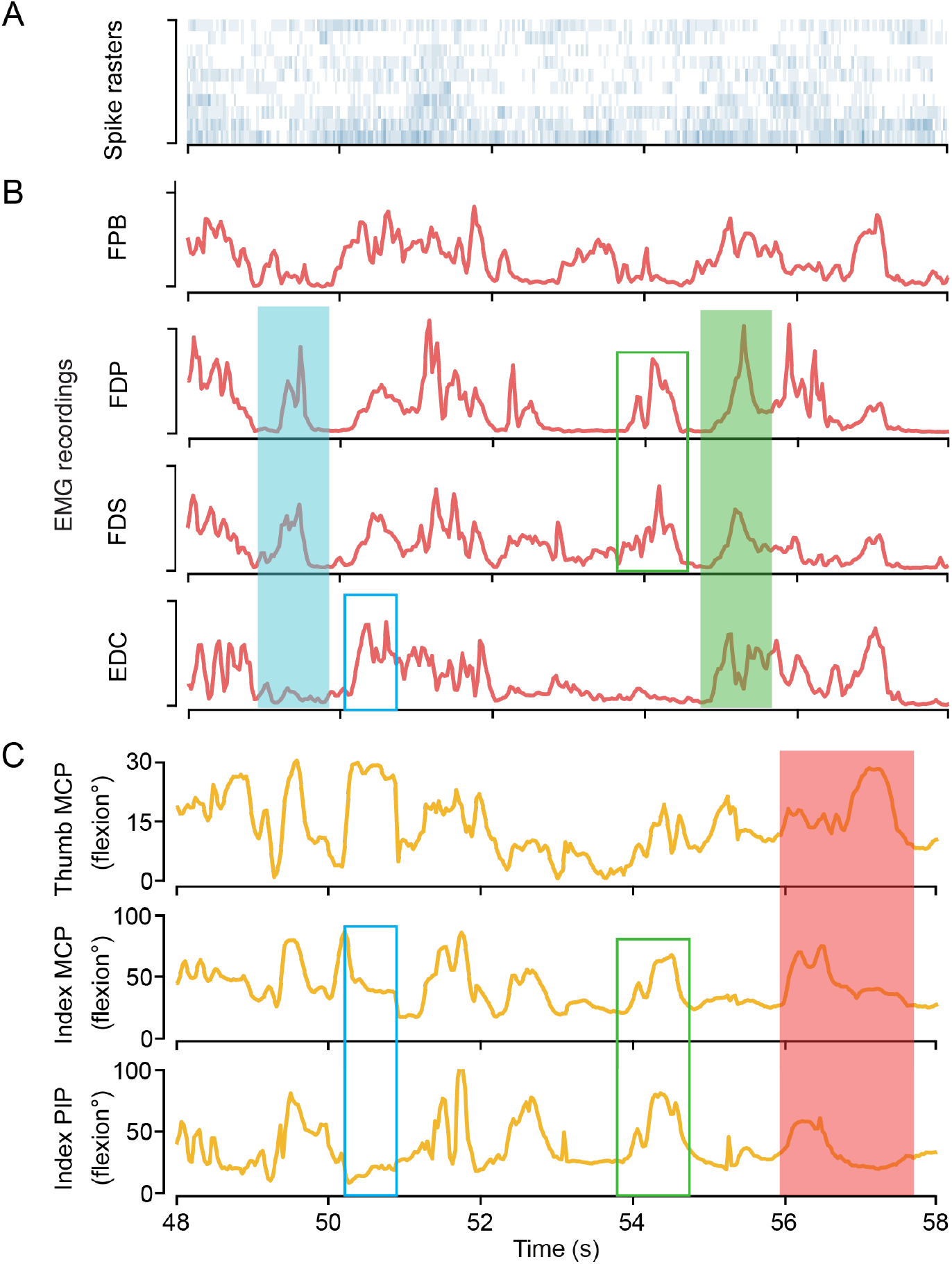
Representative eight-second segment recorded from monkey TO during the grasp task. (A) Spike rasters for the 10 most strongly modulated neurons. (B) EMG activity from four muscles involved in index finger and thumb movements: *flexor pollicis brevis* (FPB), *flexor digitorum profundus* (FDP), *flexor digitorum superficialis* (FDS), and *extensor digitorum communis* (EDC). (C) Joint angles for thumb metacarpophalangeal (MCP) flexion/extension, index MCP flexion/extension, and index proximal interphalangeal (PIP) flexion/extension. 0 deg flexion corresponds to fully extended joints. Filled and empty boxes highlight behavioral events described in the text.

Not surprisingly, during the grasp task, FDP and FDS – the finger flexors inserting on the distal and middle phalanges, respectively – were strongly correlated (cyan shading), while distinct in their activation from the intrinsic thumb flexor FPB. EDC, the common digit extensor, was at times reciprocally related to the flexors (cyan shading), and at other times coactivated (green shading). Likewise, the index MCP and PIP joints were well correlated with each other but less with the thumb MCP joint (red shading). Although the relationship between muscle activity and joint rotation is non-linear, activation of finger flexors does lead to digit flexion (paired green boxes), and activation of extensors – to extension (cyan boxes). While it is possible to see some correspondence between the EMG and position signals in these examples, their relation to M1 firing rates is far more difficult to appreciate in this qualitative view. Instead, we constructed generalized linear models (GLMs) from the EMG and kinematic signals to neural firing rate.

### Continuous encoding models of M1 firing rate

Figure 3 illustrates the qualitatively different results of EMG and kinematic encoding models for two representative neurons. Figure 3A shows the actual and predicted firing rates for a single neuron collected from monkey TO during the grasp task. While the position-based model consistently fit the firing rate quite accurately, the joint velocity inputs were much less accurate. To compare these predictions quantitatively, we computed the pseudo-R^2^ (pR^2^) ^17^ between the recorded and modeled signals. The visual impression of prediction accuracy for these short segments was consistent with the pR^2^ values for the entire two-minute recording from which they were taken (angular position pR^2^ = 0.39, angular velocity pR^2^ = 0.05). The GLM with EMG inputs had essentially the same pR^2^ as that of position (pR^2^ = 0.41).

**Figure 3:**
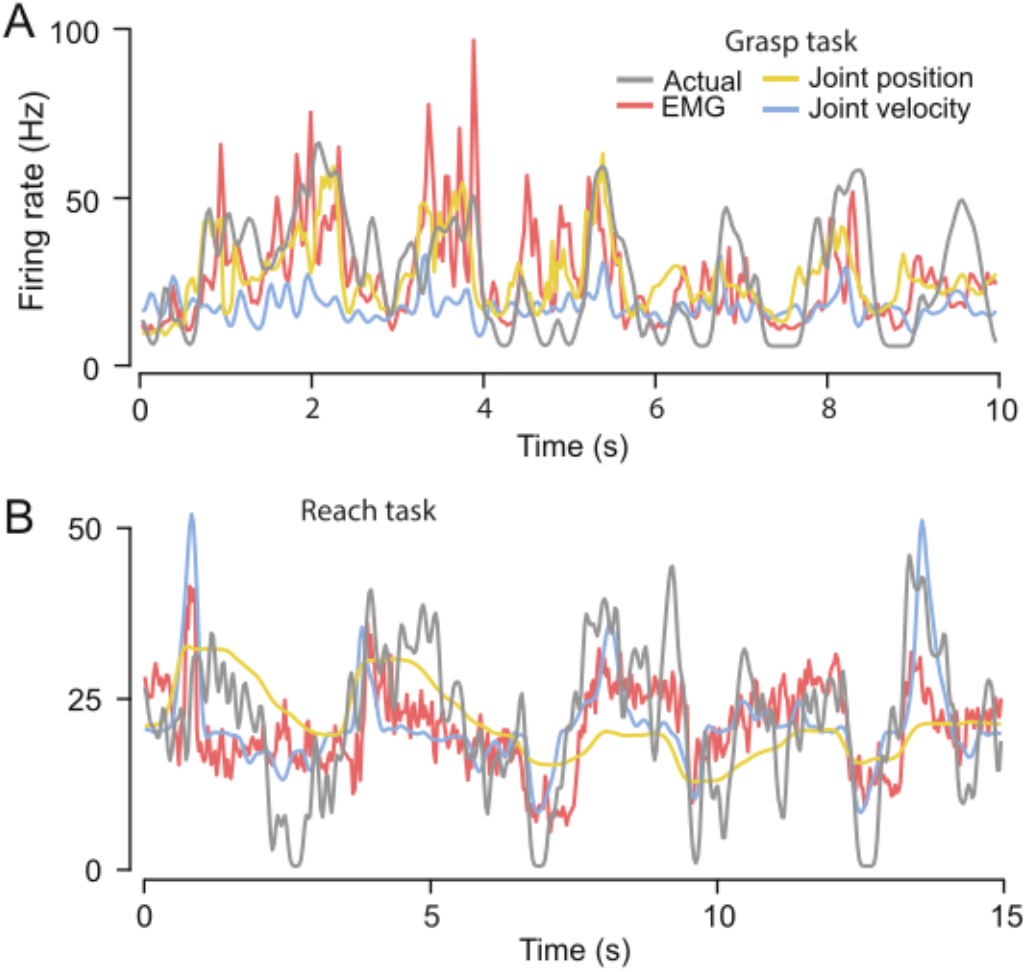
Firing rate predictions from GLM encoding models using different input modalities. (A) Actual and predicted firing rates for an example neuron from Monkey TO during the grasp task. Grey lines indicate actual firing rate. Coloured lines show predictions from encoding models using EMG (red), joint angular position (yellow), and joint angular velocity (blue) as inputs. (B) Actual and predicted firing rates for an example neuron from Monkey AR during the reach task, with the same colour conventions.

Figure 3B shows the analogous result for a single neuron from monkey AR during the center-out reach task. Here, the kinematic encoding results were quite different. Unlike the grasp task, for which position provided an accurate estimate of firing rate, in this case, pR^2^ for position was only 0.13, while for velocity, it was 0.24. The accuracy of the EMG-based model was 0.28, again similar to the better of the two kinematic models.

Whatever M1 may encode, the descending signals must first activate muscles, resulting in EMG, force production, and ultimately movement. Therefore, we expect to find different delays between neural activity and each class of input signal. We identified optimal lags for the three input classes: EMG signals, position (or angular position, as appropriate), and velocity (or angular velocity). We systematically tested lags ranging from −1000 ms to +1000 ms in 33 ms increments for all predictors of the same class. We evaluated the accuracy of the models by computing the pR^2^ between actual and predicted firing rates. Figure 4 contains histograms of the optimal lags across neurons for the different monkeys and input classes. We defined the optimal lag for each signal class and monkey as the mode of each of these distributions.

**Figure 4:**
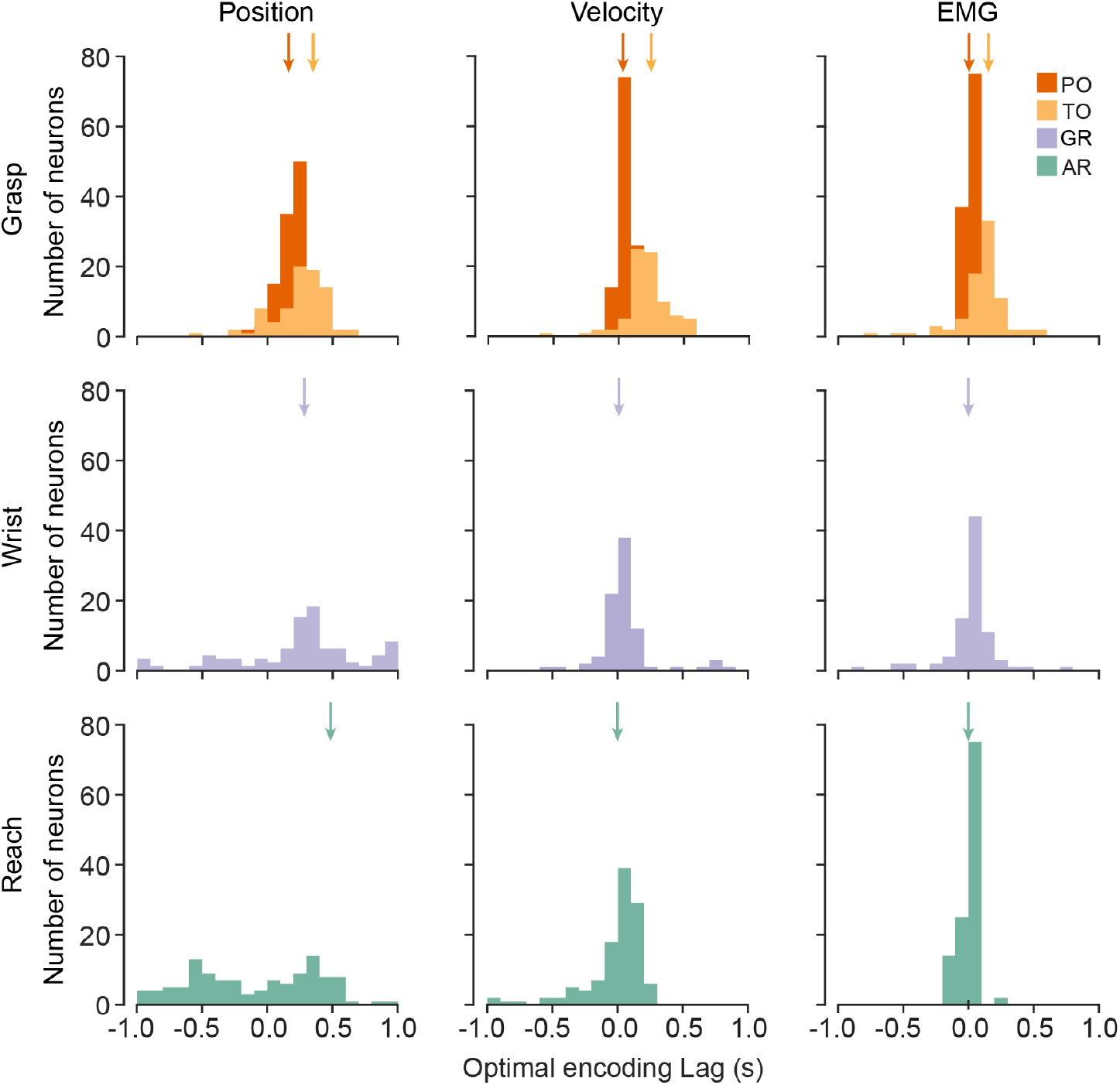
Optimal lag distributions for each signal type and task. Distributions of optimal input–output lags across neurons for each encoding model input class. We defined the optimal lag for a given neuron as the value that maximized pseudo-R^2^ in its GLM encoding model. The upper row shows the grasp task lags (monkeys PO and TO) for joint angles, joint velocities, and EMG signals. The middle row shows the analogous results for the wrist task (monkey GR). The bottom row shows the reach task (monkey AR). From these results, we defined the optimal lag for each monkey/task combination as the mode of each distribution (coloured vertical arrows) These modes and interquartile ranges (IQR) are listed in the text.

Distinct lag profiles emerged across signal types. Hand position and joint angle lags (left column) were distributed rather broadly, particularly for the reach and wrist tasks. The mode for all tasks was at or above 200ms (PO: 200 [IQR: 133–233] ms, TO: 367 [133–367] ms, AR: 467 [-533– 333] ms, GR: 333 [150–467] ms). Velocity lags (middle column) were shorter and more tightly distributed than position (PO: 67 [33–67] ms, TO: 233 [133–300] ms, AR: 0 [-67–133] ms, GR: 33 [-33–67] ms). EMG lags (right column) were even shorter and slightly more tightly distributed (PO: 0 [-33–33] ms, TO: 100 [67–167] ms, AR: 0 [-83–67] ms, GR: 0 [-33–67] ms) than velocity. These results reinforce the importance of incorporating appropriate delays for the inputs to maximize the accuracy of the GLM encoding models. In subsequent analyses, we used the optimal lag for all neurons for a given task, signal type, and monkey.

### Generalized Linear Model accuracy varies with different classes of inputs

We then went on to compare the accuracy for different input classes for each task, summarized in Figure 5 with tasks in rows and comparison of a particular pair of input variables in columns. The first column compares velocity and position encoding models for all three tasks for each neuron. For the grasp task, the position was encoded much more strongly than was velocity (Figure 5A), consistent with the single example shown in Figure 3A. Position-based models outperformed velocity for virtually every neuron from both monkeys, with the difference across neurons highly significant (210 neurons, paired Wilcoxon signed-rank test, p < 0.0001, rank-biserial correlation coefficient r_rb_ = 0.99). In contrast, kinematic encoding models for the reach task (Figure 5G) were weighted toward velocity inputs, with a more modest effect size (120 neurons, paired Wilcoxon signed-rank test, p = 0.0003, r_rb_ = −0.38). We confirmed these results in two additional monkeys (CH and MH) that did not have EMG implants (Extended Data Figure 1). Finally, results shown in Figure 5D for the wrist task were intermediate between those of the grasp and reach tasks with no significant difference between position and velocity encoding across neurons (n = 90 neurons, paired Wilcoxon signed-rank test, p = 0.45, r_rb_ = 0.09).

**Figure 5:**
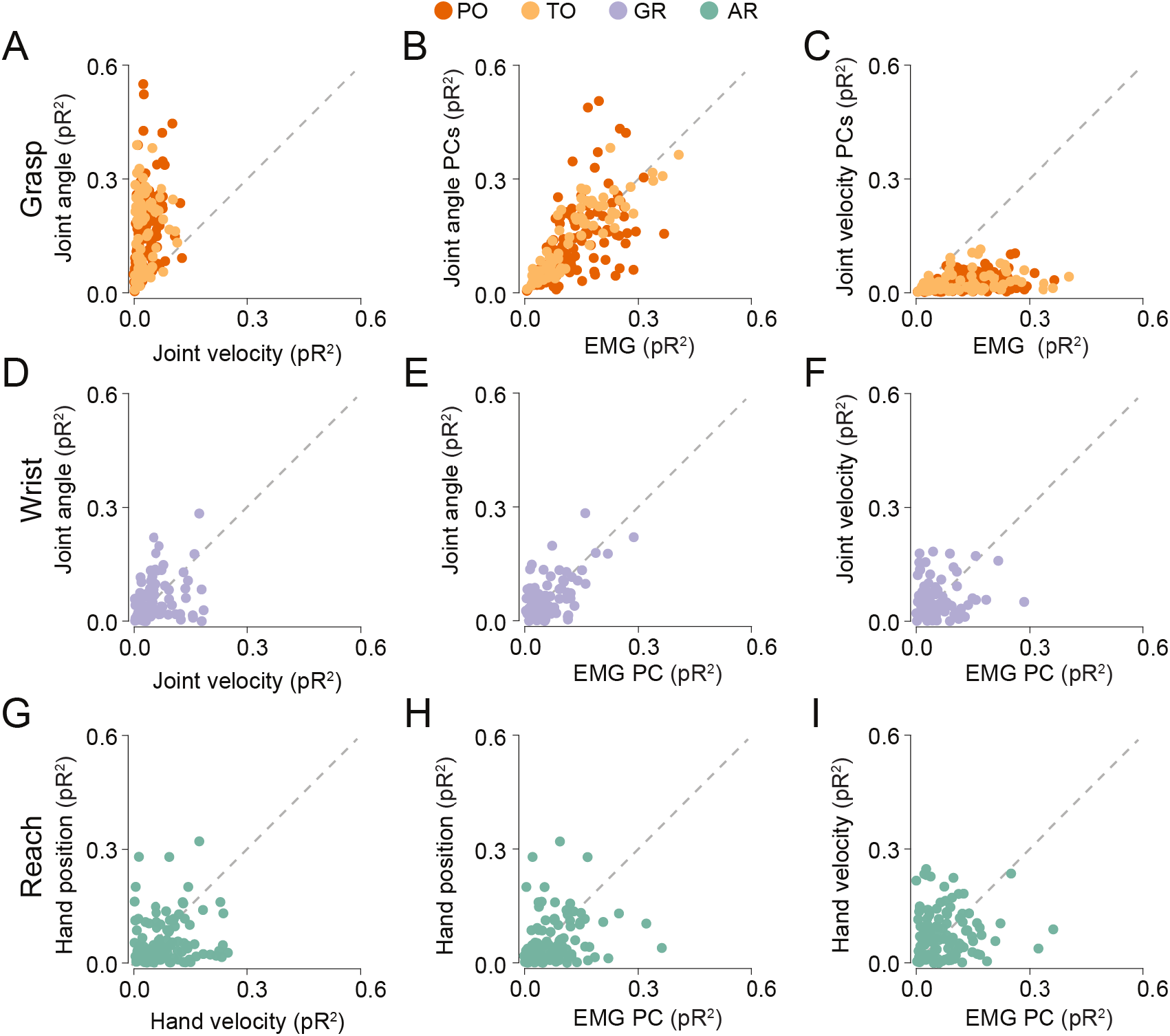
Encoding model performance across tasks and input modalities. Pairwise comparisons of pseudo-R^2^ values across neurons for different encoding model inputs, organized by task (rows) and comparison type (columns). (A–C) Grasp task: joint angles versus joint velocities (A), joint angle PCs versus EMG (B), and joint velocity PCs versus EMG (C). (D–F) Wrist task: joint angles versus joint velocities (D), joint angle versus EMG PCs (E), and joint velocity versus EMG PCs (F). (G–I) Arm reach task: hand position versus hand velocity (G), hand position versus EMG PCs (H), and hand velocity versus EMG PCs (I). Each point represents one neuron; points above the diagonal indicate better performance for the variable on the y-axis. For the grasp task, position-based models strongly outperformed velocity; for the reach task, velocity outperformed position; for the wrist task, performance was intermediate. EMG-based models matched or exceeded the best kinematic model in each task.

The middle and rightmost columns compare the kinematic and EMG encoding models. It is important to recognize that, while the comparison between the two classes of kinematic inputs is a fair one, a direct comparison between EMG and kinematic inputs is less so. This is in large part because of the difference in numbers of degrees of freedom for the two classes. In the grasp task, there were many more kinematic degrees of freedom (24, albeit certainly not all independent) than there were recorded muscles (7-17 depending on the monkey and session). During the planar reach task, the opposite was true. We used only X and Y coordinates of hand motion, but EMG activity was recorded from a larger set of muscles (9–11 depending on the session). To deal with this mismatch, for any given comparison we computed principal components for the larger class of inputs and retained only the number of components equal to that of the smaller input class.

There was no significant difference between joint angle PCs and EMG model performance for the grasp task (Figure 5B, 210 neurons, p = 0.46, r_rb_ = 0.06). As anticipated, the remaining comparison between angular velocity PCs and EMG strongly favored EMG (Figure 5C; 210 neurons, p < 0.0001, r_rb_ = −0.99). This was confirmed in a single session without the forearm restraint (See methods and Extended Data Figure 2). Unlike grasp, for the reach tasks, EMG models outperformed those based on position (Figure 5H; 120 neurons, p < 0.0001, r_rb_ = −0.33) and were as accurate as models based on velocity (Figure 5I; 120 neurons, p = 0.56, r_rb_ = 0.06). Encoding performance during the wrist task was more varied across neurons. EMG-based models achieved comparable accuracy to both position-based (Figure 5E; 90 neurons, p = 0.39, r_rb_ = −0.10) and velocity-based models (Figure 5F; 90 neurons, p = 0.57, r_rb_ = −0.07). Overall, models encoding EMG activity performed as well as the best kinematic model for each task.

### GLM performance is consistent with differences in limb dynamics

We sought to explain how the difference in the kinematic encoding could arise despite the consistent encoding of the EMG signal. We tested whether differences in mechanical impedance between limb segments could provide the explanation, by computing impulse response functions (IRFs) between select pairs of muscles and the joints they actuate for each task.

The shape of an IRF reveals the internal dynamics of a system, specifically, how it would respond to a brief, “impulsive”, input. For example, if a system does not alter the signal much, but simply adds a transmission delay, then the response will retain the shape of the impulse at an appropriate delay. On the other hand, if it were to act as an integrator, it would transform the brief input to a step function. Figure 6A shows example IRFs computed for synthetic signals which had the same sampling rate and duration as the actual signals but containing a white noise as input. In the upper example, the output signal was simply a copy of the input, delayed by four sample points, and with added independent noise of equal variance. The narrow, delayed peak reveals the 120 ms lag, with no additional alteration. Below it is the IRF for a similar output signal, but with the further addition of an integrator. As anticipated, it has the form of a noisy step function, beginning after a short delay.

**Figure 6:**
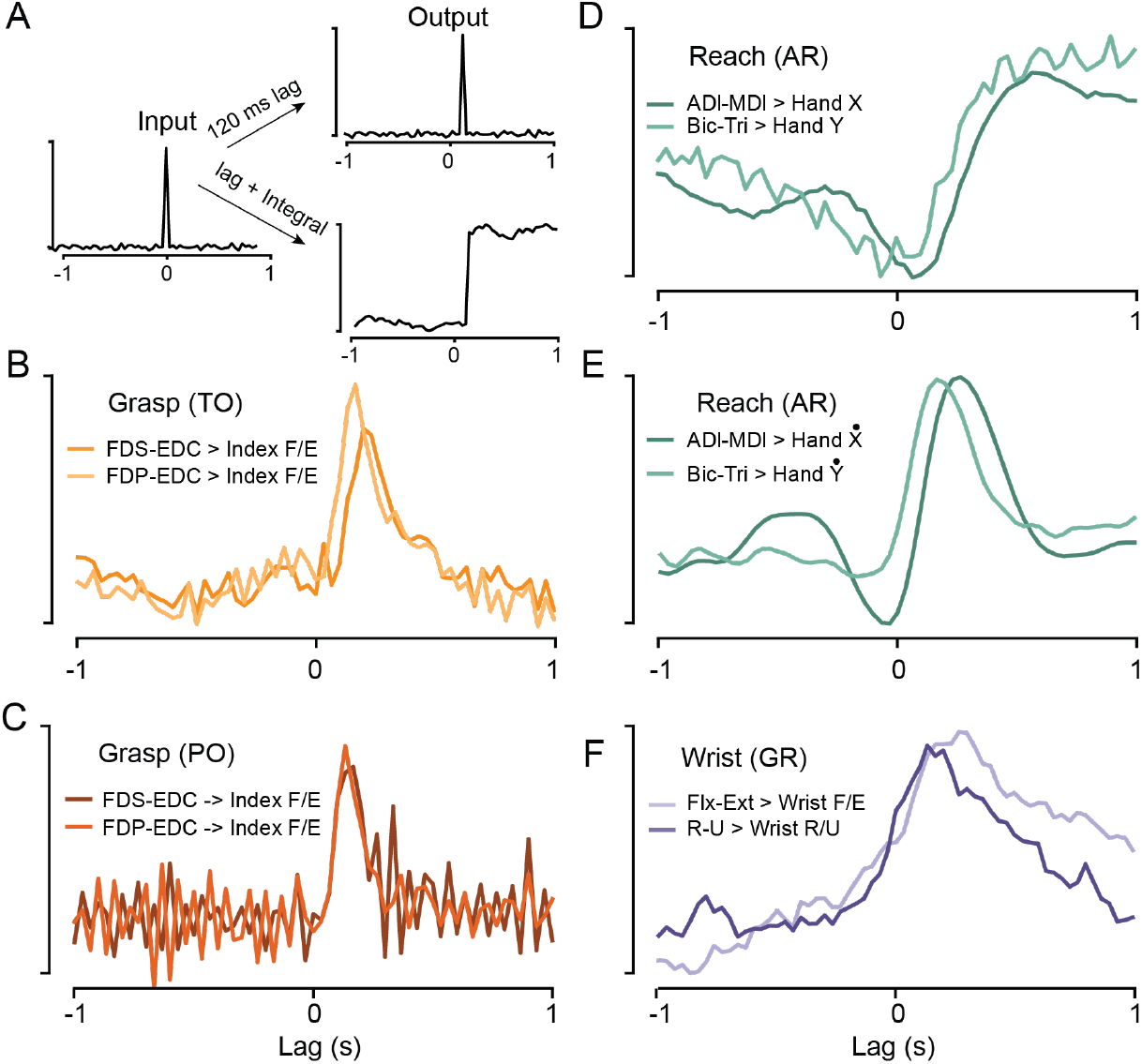
Impulse response functions (IRFs) between muscle activity and joint kinematics. (A) IRFs computed from synthetic signals, illustrating responses to additive noise and integrator dynamics. (B-F) Representative IRFs from experimental data, computed between EMG of antagonistic muscles and a single kinematic degree of freedom. (B,C) Grasp task for monkeys TO and PO, revealing transmission delays of ∼100 ms and a fast decay for index finger flexion/extension and two different pairs of muscles. (D) Reaching task for monkey AR had integrator-like dynamics with a ∼200 ms lag for two different pairs of muscles corresponding to movements along the x- and y-axes (forward-back and medial-lateral), respectively. (E) Same as D, but for velocity of x and y instead of position. (F) Monkey GR wrist task, with dynamics intermediate between hand and proximal arm movements for both radial/ulnar deviation and flexion/extension DOFs.

Figure 6B-F were computed for select signals from representative sessions for the three tasks. In each case, we considered the position of a single DOF as the output signal and the pair of muscles that actuate that DOF as the input. We subtracted one of the EMG signals from the other to form a signal that was modulated in both directions. Figure 6B shows the IRF for index finger flexion-extension during the grasp task (Monkey TO). Joint flexion is positive, and we used both EDC subtracted from FDS (FDS-EDC) and FDP-EDC as input signals. Both IRFs suggest there was some low pass-filtering which broadened the pulse, as well as a transmission delay of roughly 100ms. Although slightly noisier, the IRFs in Figure 6C for monkey PO were essentially the same, suggesting a delay but only modest dynamic transformation between these EMGs and joint angular position.

Both examples stand in sharp contrast to an IRF calculated for hand position during the reach task (monkey AR). With the arm in the sagittal plane, movements of the hand away from the body (the x-axis) were primarily associated with the Anterior (ADl) and Medial (MDl) deltoids (hence ADl-MDl), while movements across the body (y-axis) were more strongly related to biceps and triceps (Bic-Tri). In both cases, the dynamics were close to those of an integrator function with a transmission delay of approximately 200ms. These results suggest that the dynamics of the proximal limb are act to integrate the EMG signal. Consistent with this interpretation is the IRF in panel E, which was computed between the same muscles as Figure 6D, but velocity has the output signal. In this case, the shape is very much like that in panels B and C, reflecting relatively little dynamical transformation between EMG and velocity. Finally, Figure 6F shows the IRF for the wrist task (monkey GR). Because of the arrangement of wrist muscles in the forearm, we used combinations of muscles corresponding most closely to the flexion/extension (FCR+FCU-ECR-ECU), and radial/ulnar deviation (FCR+ECR-FCU-ECU) DOFs. As in the encoding results of Figure 5, the resultant wrist dynamics were intermediate between those of the hand and the arm.

We constructed a mathematical model of a single-joint system in a viscous environment with a gravity force field to identify the primary physical properties responsible for these dynamical differences (Supplementary Equations). Although the specific solutions to the system involve many parameters, the speed of decay in the IRF was inversely proportional to the mass and length of the modeled segment. A segment with little mass and short length produces an IRF with a very fast decay, essentially leaving the input signal unchanged, as in the hand. On the contrary, long, massive segments lead to slow decay in IRF, more nearly resembling the step function we observed for the arm. Thus, simply the difference in size and mass of a segment could produce the observed differences in the EMG/kinematic relationships.

## Discussion

Understanding the neural mechanisms by which M1 controls dexterous hand use remains an intriguing question in neuroscience, one which has been the topic of many experimental and theoretical studies. In this study, we used chronically implanted intramuscular electrodes in the arm and hand to directly compare kinematic and EMG encoding models of cortical activity during reaching, wrist movement, and grasping. We replicated the observations of earlier studies showing strong correlation between M1 and hand velocity during reaching ^3-7^, and the seemingly contradictory observation of equally strong correlation between M1 and joint angular position during grasping ^8-10^. Critically, we found that across all three motor behaviors we studied, EMG encoding models explained the activity of M1 neurons as well as the best kinematic models, leading us to ask whether the variable results with kinematic models may simply be explained by the different dynamics of the arm and hand.

Decades of earlier experiments and analyses have been devoted to comparing neuromodulation with different variables describing movement. A large body of work examining reaching movements as well as wrist movement and grasp (typically in separate studies), has led to conflicting hypotheses regarding whether M1 primarily encodes kinematic variables, such as position or velocity, or whether it more directly represents muscle activation patterns ^1,3,18-21^. This question has gained renewed attention following the report of a difference in the relation between M1 activity and kinematics during grasping and reaching. In contrast to the classic observation of hand-velocity tuning during reaching ^3^, M1 during grasp correlated much more strongly with digit joint angular position ^8^. Several explanations were proposed for this apparent discrepancy, including the possibility that proximal and distal limb movements are represented through fundamentally different cortical mechanisms ^8^. The greater tendency of hand area corticospinal neurons to project directly to motoneurons compared to that of the proximal arm ^22^ lends credence to this interpretation.

An important series of experiments examined the dynamics of M1 activity during ramp and hold wrist movements against both elastic and isometric loads. Even for these very simple movements, the authors reported a wide variety of activation patterns across 135 corticomotoneurons, that included all combinations of phasic bursts of activity, tonically maintained activity, and ramping activity ^23^. The authors speculated these disparate dynamical components were assembled to generate the corresponding components of the muscle activation. Other studies have suggested that the contribution a given neuron makes to its target muscle activity may vary with that muscle’s function in a particular task, differing for power and precision grasp ^24^, or with the orientation of the forearm ^25^. Some of these unexpected, changing relationships between M1 activity and movement may be related to, or at least analogous to, similar counterintuitive relationships between EMG and kinematics due to complex biomechanics. For example, triceps is modulated during pronation–supination movements despite having no mechanical role in producing those torques, in order to counteract the strong biceps flexion torque ^26^. Likewise, wrist extensors are activated during grasp, to counteract the wrist flexion torque produced by the extrinsic finger flexor tendons which cross the wrist ^27^.

The published literature often describes the proximal arm as dominated by inertial and intersegmental interaction forces ^14,28^. On the other hand, hand-related forces are more related to passive elastic forces and the stiffness properties of active muscle ^29 30^. Elastic-like responses in cortical neurons have also been ascribed to the length-tension properties of muscles ^31^. We wanted to examine if the difference in the kinematic encoding could be explained by limb mechanics. To pursue this possibility, we conducted a simple systems identification analysis, using impulse response functions (IRFs) ^32,33^ to characterize the dynamical transformation between muscle activity and kinematics. In our data, EMG dynamics were more strongly associated with velocity-related components for proximal arm movements and with position-related variables for hand movements. Although the relationship between EMG and kinematic variables is complex and hard to model precisely^34^, we have also shown, using a model of a simple mass/damping system, that the dynamical differences can be explained largely by the difference in the mass and length of coupled segments (Supplementary Equations). This combination of empirical, modeling and theoretical results all point to system in which a simple dynamical relation between M1 neurons and muscles is maintained across tasks.

This discovery may also have practical implications for brain computer interfaces (BCIs), nearly all of which, whether for the arm ^35,36^or hand^37^, are based on a mapping to kinematics. As such, they would be expected to perform suboptimally as dynamical conditions vary, requiring the user to alter their strategy for different movement contexts. By incorporating muscle-based, biomechanically-informed models, BCIs might achieve more robust generalization across a broader repertoire of natural movements.

Even though we have argued that the activity of motor cortical neurons is more consistently related to muscle activity than it is to kinematics, we do not claim that it is the sole determinant of muscle activity. Obviously, multiple neurons, many with different dynamics, contribute to the activity of any given muscle. These neurons are not confined to the motor cortex but are also found in brainstem motor areas and the spinal cord. It is likely these different sources of input to spinal motor neurons make functionally different contributions. There is evidence that neurons in the magnocellular red nucleus related to the arm preferentially control distal extensors ^38,39^. In contrast, the reticular formation appears to be more closely related to elbow and forearm flexors ^40,41^. Nonetheless, the organization of the descending inputs would be simplified, to the extent that these multiple convergent sources all contribute components of muscle activity.

It must be noted that other studies have shown that M1 neurons tend to be more phasic during wrist movement than many EMGs ^23^. There is also some evidence from spinal recordings during wrist movement, that spinal interneurons do effect some amount of dynamical transformation of the descending inputs to the cord, transforming relatively phasic commands into longer-lasting, sustained control signals ^11^. Another recent study combining a variety of human and monkey data with simulation, even posits a spinal (or at least, sub-cortical) integrator of descending cortical activity ^13^ analogous to that of eye movements ^42,43^. Although a fascinating possibility, it is difficult to see how (unlike the case for the eye) an integrator could serve to transform an arm movement command into an appropriate postural command, given the constantly varying dynamics of the upper limb.

In summary, the mechanical differences between arm, wrist, and hand likely contribute substantially to the observed task-dependent differences in kinematic encoding without the need to invoke distinct cortical mechanisms. Instead, muscle-related cortical signals would naturally appear position-tuned when filtered through the elastic-dominated dynamics of the hand, and velocity-tuned when filtered through the inertial- and intersegmental-dynamics of the arm.

## Methods

### Animal and Surgery

All surgical and experimental procedures were approved by the Institutional Animal Care and Use Committee (IACUC) of Northwestern University. Data were collected from six male macaque monkeys. Most monkeys were implanted with a 96-electrode array with 1.5 mm electrode length (Blackrock Microsystems, Salt Lake City, UT) in either the hand or arm area of the primary motor cortex (M1). Monkey TO had a 128-electrode array. During surgery, M1 was identified using visual landmarks and bipolar cortical surface stimulation to evoke twitches of distal or proximal muscles. The arrays were then inserted pneumatically.

In separate surgical procedures, four of the monkeys (PO, TO, AR, GR) were also implanted with intramuscular electromyographic (EMG) leads in the contralateral arm, forearm and hand muscles. Electrode location was verified intraoperatively by evoking muscle contractions through stimulation. Two of the reach-task monkeys (CH and MH) had no implanted EMG electrodes, and were used to confirm the kinematic model observations of monkey AR.

### Behavioral Tasks

#### Grasp

Monkeys PO and TO were seated in a primate chair with the arm supported to restrict proximal limb movements. A custom plastic support maintained the elbow at a fixed 90° angle and rigidly constrained the upper arm and forearm, effectively preventing reaching movements and limiting voluntary motion to the wrist, hand, and digits. This configuration allowed the monkey to perform grasp-and-release movements without translating the arm in space. We conducted a single session with monkey PO, without the forearm restraint, allowing free movement of the forearm as well as the wrist and fingers. As results in the two conditions were similar, but motion tracking was more accurate with the forearm constraint, we adopted it for subsequent experiments. In each session, the trainer presented various objects mounted on the end of a thin rod, pulling them away just before contact to minimize tactile effects. These objects included a 2cm sphere, a large cylinder with length 5 cm and a diameter 2 cm, and a small (4 cm × 1 cm) cylinder (Figure 1A). The cylinders could be mounted either parallel or perpendicular to the rod. The trajectory and speed of the objects were varied across trials to elicit diverse, uncorrelated finger movements and grasps. Water or juice rewards were delivered periodically to reward correct task engagement.

#### Wrist

We trained one male rhesus monkey (monkey GR) to sit in the primate chair with the forearm restrained, grasp a wrist-operated manipulandum, and make wrist flexion/extension and radial/ulnar deviation movements to control the movement of a cursor (Figure 1B). A trial began when the monkey moved the cursor to a central target, requiring the wrist to be placed in a neutral position with the forearm and hand aligned. After a hold time of 500ms, a randomly chosen outer target appeared. To complete a trial successfully and receive a reward, the monkey had to reach this target within five seconds and hold for 500ms.

#### Reach

Monkeys AR, CH and MH were trained to sit in a primate chair and grasp the handle of a two-link planar manipulandum that controlled a 1cm diameter cursor on a screen within a 20×20 cm workspace (Figure 1C). The monkeys performed a standard center-out (CO) task, initiating a trial by moving to the center target. After a hold period of 500-1500 ms, a 2 cm target was randomly displayed in one of eight regularly spaced outer positions at a radial distance of 8 cm, followed by a variable delay period of 500-1500 ms before an auditory go cue. The monkeys were required to reach the target within one second and hold for 500 ms to receive a liquid reward.

### Data acquisition and pre-processing

#### Neural Data

Neural activity was recorded using a Cerebus system (Blackrock Neurotech) and amplified with a Cereplex-E headstage. Spike sorting was performed manually with Offline Sorter (Plexon Inc.) to isolate putative single units. Neural data were binned at 30 Hz to match the motion capture sampling rate (below) and smoothed with a Gaussian kernel (σ = 50 ms). For subsequent analyses, the 30 neurons with the highest modulation depth (difference between 90th and 5th percentile of firing rate, computed with 20 ms bins) were selected.

#### EMG Data

EMG signals were collected using a multi-channel amplifier and a wired transmitter. The recorded EMG signals were bandpass filtered (4-pole Butterworth, 50-500 Hz). To extract the envelopes, signals were subsequently rectified and low-pass filtered (4-pole Butterworth, 10 Hz) and then subsampled to 30 Hz to correspond to the motion capture kinematic data.

#### Kinematic Data (grasp task)

Hand movements were recorded using four ImagingSource DFK 37BUX265 cameras at 30 frames per second with 1280 *χ* 1024 resolution mounted on a metal frame surrounding the workspace. A series of processing steps were implemented to extract time-varying joint angles of the hand from the video.

More specifically, we first used Jarvis MoCap (https://github.com/JARVIS-MoCap) markerless motion tracking system to track the positions of 24 landmarks on the wrist and fingers. We labeled 1750 frames per animal to train the machine vision networks. The quality of the tracking was assessed through visual inspection of predicted joint positions superimposed on the video. Missing values in the data (e.g., when a digit was obscured) were filled using linear interpolation. Outliers were identified as elements deviating more than three times from the local median within a moving window of 100 samples, and these outliers were replaced by the nearest valid value. The data were then smoothed using a Butterworth low-pass filter of the fourth order with a cut-off frequency of 7 Hz.

After obtaining 3D positions of landmarks, we computed time-varying joint angles using the inverse kinematic tool in OpenSim ^44^, and a musculoskeletal model (https://github.com/nishbo/ms_arm_and_hand) scaled to each subject. The joint angles were filtered with a 7 Hz fourth-order Butterworth low-pass filter. We computed joint velocities by taking the derivative of the joint angles followed by a 5 Hz fourth-order Butterworth low-pass filter.

#### Kinematic Data (reach task)

Two-dimensional hand position was computed from rotary encoders mounted on the joints of the planar manipulandum. Position and velocity signals were then low-pass filtered using a fourth-order Butterworth filter with cut-off frequencies of 7 Hz and 5 Hz, respectively.

#### Kinematic Data (wrist movement task)

Wrist kinematics were measured directly from joint angle encoders, capturing flexion/extension and radial/ulnar deviation. Position and velocity signals were low-pass filtered with a fourth-order Butterworth filter at 7 Hz and 5 Hz, respectively.

### Calculation of encoding models

To investigate the type of information encoded by M1 neurons, we modeled neural activity with respect to the movement covariates using Poisson generalized linear models (GLMs) ^45^. Specifically, neural activity *f* was assumed to follow a Poisson distribution with rate parameter λ, which depends on the behavioral covariates X through a log-linear relationship:

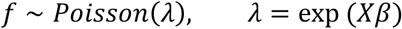

Here, *f* is a *T × N* matrix of spike counts or average firing rates across *T* time points and *N* neurons. *X* is a *T χ P* matrix of behavioral covariates—including kinematic or EMG signals—used to predict neural responses, and *β* is a *P × N* matrix of model coefficients. The exponential function ensures that the predicted firing rates are strictly positive.

GLM parameters *β* were estimated via maximum-likelihood estimation. Using the fitted models, we predicted firing rates on held-out data:

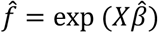

We used pseudo-R^2^ to assess model prediction accuracy by comparing the deviance (or log-likelihood) of the full model against a null model that predicts only the overall mean ^17,46^. Values range from negative (worse than null) to 1 (perfect fit), with 0 indicating no improvement over the null model. Unlike traditional R^2^, a pseudo-R^2^ of above 0.2 is considered a good fit in logistic regression contexts ^47^.

To account for the physiological delay between firing rates and covariates, we found the lag between inputs and output which maximized pR^2^, testing lags between ±1000 in 33 ms increments (corresponding to the video frame rate). We considered lags for which model input data (EMG/kinematics) followed M1 firing rate to be positive, since the direction of the encoding models was opposite causality. Optimal lag parameters were determined by finding the mode of lag distributions for each type of input signal and monkey.

We tested encoding models with 11 different sets of inputs, shown in Extended Data Table 3. The models differed in the type of input features considered, including single-actuators (such as hand positions or joint angles, and hand or joint velocities) and muscle activity, as well as low-dimensional representations derived from the top principal components of joint angles, velocities, or EMGs. All analyses were conducted with the optimal delay between input and output data.

### Statistical analyses

For uniformity in the optimal lag analysis, we used the 30 Hz frame rate of the grasp task video motion capture system as the sampling rate for all tasks. We computed the mode and interquartile range of each distribution in Figure 4 on this basis.

To evaluate encoding model performance, we performed pairwise comparisons of pR^2^ values between the models using different input sets. For each neuron, pR^2^ values were averaged across cross-validation folds. Statistical comparisons were then performed across neurons pooled across all sessions for a given task.

Prior to hypothesis testing, we assessed the normality of the pR^2^ distributions using the Anderson-Darling test. If either distribution in a pairwise comparison deviated significantly from normality (p < 0.05), we used the non-parametric Wilcoxon signed-rank test for paired samples. Otherwise, we used a paired t-test. Statistical significance was assessed at α = 0.05.

To quantify the magnitude of differences between models, we computed the rank-biserial correlation coefficient (r_rb_) as a measure of effect size for the Wilcoxon signed-rank test. The rank-biserial correlation was calculated as:

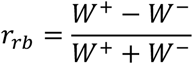

where W^+^ and W^−^ represent the sum of ranks for positive and negative differences, respectively. This metric ranges from −1 to +1, with positive values indicating that the first model outperformed the second, and negative values indicating the opposite. Effect sizes were interpreted according to established thresholds: |r| < 0.1 as negligible, 0.1 ≤‐‐ |r| < 0.3 as small, 0.3 ≤‐‐ |r| < 0.5 as medium, and |r| ≥ 0.5 as large.

### Calculation of impulse responses

Given measured input and output signals, it is possible to compute IRFs by deconvolving the input (removing the effect of its dynamics) from the output. We used custom python code to compute two-sided IRFs in the frequency domain, using EMG signals as the input and position (or velocity) as the output.

## Author Contributions

***Conception: XM, LEM***

***Experiments: CC, XM, EG***

***Analysis: CC, XM, FR, ARS, LEM***

***First draft: CC, ARS, LEM***

***Approved Final draft: CC, XM, FR, ARS, EG, LEM***

## Supplementary Equations

Consider a general oscillator system describing a single damped joint with external input in the form of torque, an external force field (e.g., gravity), and mass m at length L:

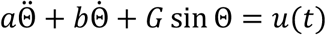

Where *a* = *mL*^2^, b = viscosity of the joint, *G* = *mgL* in case of moving in gravity. The system can be linearized around a point:

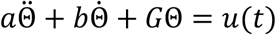

This equation in Laplace space, assuming starting position and velocity 0:

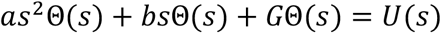

Dividing by Θ(*s*) and taking reciprocal, one can find the transfer function from *U*(*s*) to Θ(*s*):

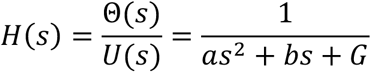

Define values for simplicity:

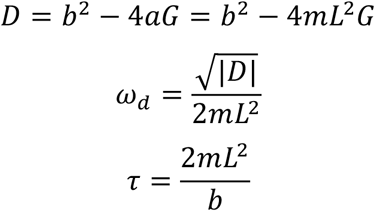

The inverse Laplace transform of the transfer function yields the impulse response of the system. There are two cases:

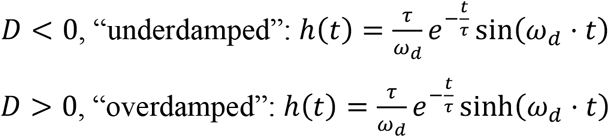

The exact relationship between the damping and the mass, length, gravity is hard to estimate for a specific body part, especially since the damping is not well known and can depend on muscle coactivation. Therefore, we are not going to claim whether the system is under or overdamped, or the exact period of the oscillation. However, the decay factor *τ* depends inversely on mass and segment (and muscle) length. Thus, a larger, more massive system like the arm will have an impulse-response that decays much slower than that of the hand.

**Extended Data Figure 1.**
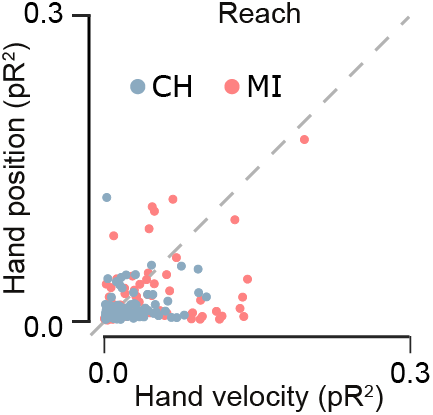
Kinematic encoding model performance for the reach task in two additional monkeys. Pairwise comparison of pseudo-R^2^ values for hand position versus hand velocity encoding models, shown for each neuron recorded from monkeys MI and CH during a center-out reach task. Each point represents one neuron; points above the diagonal indicate better performance for hand velocity, while points below indicate better performance for hand position. As in Figure 5G (main text), velocity-based models outperform position-based models for the majority of neurons, consistent with the pattern observed in the primary cohort. Note that EMG data were not collected for these two animals, precluding EMG-based comparisons.

**Extended Data Figure 2.**
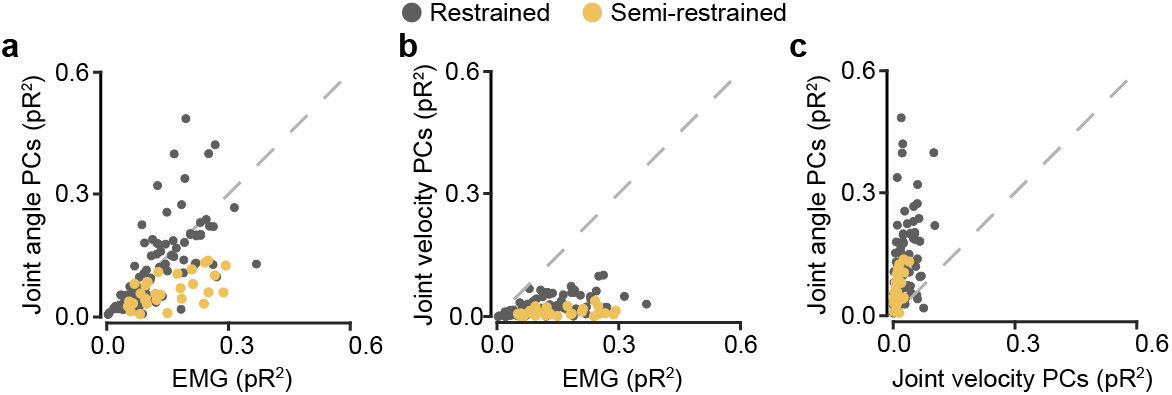
Encoding model performance for the grasp task under fully restrained and semi-restrained conditions. Pairwise comparisons of pseudo-R^2^ values across neurons for joint angle PCs versus EMG (A), joint velocity PCs versus EMG (B), and joint angle PCs versus joint velocity PCs (C). Dark grey points correspond to the fully restrained condition (monkeys PO and TO); yellow points to the single semi-restrained session (monkey PO), in which the forearm was free to move. Semi-restrained neurons showed slightly lower overall encoding performance, potentially reflecting the additional movement variability introduced by freeing the forearm.

**Extended Data Table 1.** Summary of subjects and recording parameters.

| Monkey | Implant side | Task | EMG |
| --- | --- | --- | --- |
| PO | Hand area, M1 (right hemisphere) | Freeform grasp | Left arm/hand |
| TO | Hand area, M1 (left hemisphere) | Freeform grasp | Right arm/hand |
| GR | Hand area, M1 (right hemisphere) | Center-out wrist movement | Left arm/hand |
| AR | Arm area, M1 (left hemisphere) | Center-out reach | Right arm |
| CH | Arm area, M1 (left hemisphere) | Center-out reach | No implants |
| MH | Arm area, M1 (right hemisphere) | Center-out reach | No implants |

For each monkey, the implanted hemisphere, behavioral task, and availability of intramuscular EMG recordings are listed. Arrays were 96-electrode Utah arrays (Blackrock Microsystems) except for monkey TO, who received a 128-electrode array. M1 implant location (hand or arm area) was determined using visual landmarks and intraoperative cortical surface stimulation. EMG electrodes were implanted in muscles of the contralateral arm and hand; monkeys CH and MH had no EMG implants and were used exclusively to confirm kinematic encoding results.

**Extended Data Table 2.** Implanted muscles by monkey.

| Extended Data Table 2 \| Implanted muscles by monkey |  |  |  |  |
| --- | --- | --- | --- | --- |
| Muscles | TO | PO | GR | AR |
| 1<sup>st</sup> lumbrical | X | X |  |  |
| Flexor pollicis brevis | X | X | X |  |
| Abductor pollicis brevis | X | X | X |  |
| 1<sup>st</sup> dorsal interosseous | X | X | X |  |
| 3<sup>rd</sup> dorsal interosseous | X |  |  |  |
| Flexor carpi radialis | X | X | X | X |
| Flexor carpi ulnaris | X | X | X |  |
| Flexor digitorum profundus | X | X |  |  |
| Flexor digitorum superficialis | X | X |  |  |
| Extensor carpi radialis | X | X | X | X |
| Extensor carpi ulnaris | X | X | X |  |
| Extensor digitorum communis | X | X | X |  |
| Pronator teres | X |  | X |  |
| Supinator | X |  | X |  |
| Brachioradialis |  |  |  | X |
| Biceps | X |  |  | X |
| Triceps | X |  |  | X |
| Pectoralis clavicular |  |  |  | X |
| Latissimus dorsi |  |  |  | X |
| Middle deltoid |  |  |  | X |
| Anterior deltoid |  |  |  | X |
| Checkmarks indicate muscles from which intramuscular EMG was recorded in each monkey. Muscles are grouped from distal (intrinsic hand) to proximal. Monkeys CH and MH had no EMG implants and are omitted. |  |  |  |  |

**Extended Data Table 3.** Encoding model inputs by task.

| Extended Data Table 3 \| Encoding model inputs by task |  |  |
| --- | --- | --- |
| Task | Model input features | Number of inputs |
| Freeform grasp | Joint angles | 24 |
|  | Joint velocities | 24 |
|  | EMGs | 7-17 |
|  | Joint angle PCs | 7-17 |
|  | Joint velocity PCs | 7-17 |
| Center-out wrist movement | Joint angles | 2 |
|  | Joint velocities | 2 |
|  | EMG PCs | 2 |
| Center-out reach | Hand positions | 2 |
|  | Hand velocities | 2 |
|  | EMG PCs | 2 |
| For pairwise comparisons between kinematic and EMG models, principal components of the larger input class were computed and truncated to match the dimensionality of the smaller class (see Methods). For the freeform grasp task, the number of EMG inputs ranged from 7 to 17, depending on the number of channels with an adequate signal-to-noise ratio in each session. Accordingly, for those sessions, we matched the number of joint position and velocity principal components to the number of the EMG inputs. |  |  |

## References

1 Evarts, E. V. Relation of pyramidal tract activity to force exerted during voluntary movement. J. Neurophysiol. 31, 14–27 (1968).

2 Humphrey, D. R., Schmidt, E. M. & Thompson, W. D. Predicting measures of motor performance from multiple cortical spike trains. Science 170, 758–761 (1970).

3 Georgopoulos, A. P., Kalaska, J. F., Caminiti, R. & Massey, J. T. On the relations between the direction of two-dimensional arm movements and cell discharge in primate motor cortex. J. Neurosci. 2, 1527–1537 (1982).

4 Wang, W., Chan, S. S., Heldman, D. A. & Moran, D. W. Motor Cortical Representation of Position and Velocity During Reaching. Journal of neurophysiology 97, 4258–4270 (2007). 10.1152/jn.01180.2006

5 Ashe, J. & Georgopoulos, A. P. Movement parameters and neural activity in motor cortex and area 5. Cerebral Cortex 4, 590–600 (1994).

6 Schwartz, A. B. Direct cortical representation of drawing. Science 265, 540 –542 (1994).

7 Moran, D. W. & Schwartz, A. B. Motor cortical representation of speed and direction during reaching. J Neurophysiol 82, 2676–2692 (1999).

8 Goodman, J. M. et al. Postural Representations of the Hand in the Primate Sensorimotor Cortex. Neuron 104, 1000-1009.e1007 (2019). 10.1016/j.neuron.2019.09.004

9 Irwin, Z. T. et al. Neural control of finger movement via intracortical brain–machine interface. Journal of neural engineering 14, 066004 (2017).

10 Okorokova, E. V., Goodman, J. M., Hatsopoulos, N. G. & Bensmaia, S. J. Decoding hand kinematics from population responses in sensorimotor cortex during grasping. Journal of neural engineering 17, 046035 (2020). 10.1088/1741-2552/ab95ea

11 Shalit, U., Zinger, N., Joshua, M. & Prut, Y. Descending systems translate transient cortical commands into a sustained muscle activation signal. Cerebral Cortex 22, 1904–1914 (2012).

12 Takei, T. & Seki, K. Spinal Premotor Interneurons Mediate Dynamic and Static Motor Commands for Precision Grip in Monkeys. The Journal of Neuroscience 33, 8850–8860 (2013). 10.1523/jneurosci.4032-12.2013

13 Albert, S. T. et al. Postural control of arm and fingers through integration of movement commands. eLife 9, e52507 (2020). 10.7554/eLife.52507

14 Hollerbach, J. M. & Flash, T. Dynamic interactions between limb segments during planar arm movement. Bio. Cybernetics 44, 67–77 (1982).

15 Fromm, C. Changes of steady state activity in motor cortex consistent with the length-tension relation of muscle. Pflugers Arch 398, 318–323 (1983).

16 Zajac, F. E. Muscle and tendon: Properties, models, scaling, and application to biomechanics and motor control. Crit. Rev. Biomed. Eng. 17, 359–411 (1989).

17 Cameron, A. C. & Windmeijer, F. A. R-squared measures for count data regression models with applications to health-care utilization. Journal of Business & Economic Statistics 14, 209–220 (1996).

18 Kalaska, J. F., Cohen, D. A. D., Hyde, M. L. & Prud’homme, M. A comparison of movement direction-related versus load direction-related activity in primate motor cortex, using a two-dimensional reaching task. J. Neuroscience 9, 2080–2102 (1989).

19 Kakei, S., Hoffman, D. S. & Strick, P. L. Muscle and movement representations in the primary motor cortex. Science 285, 2136–2139 (1999).

20 Bennett, K. M. & Lemon, R. N. Corticomotoneuronal contribution to the fractionation of muscle activity during precision grip in the monkey. J Neurophysiol 75, 1826–1842 (1996).

21 Brochier, T., Spinks, R. L., Umilta, M. A. & Lemon, R. N. Patterns of muscle activity underlying object-specific grasp by the macaque monkey. J Neurophysiol 92, 1770–1782 (2004). 10.1152/jn.00976.200300976.2003 [pii]

22 Dum, R. P. & Strick, P. L. Motor areas in the frontal lobe of the primate. Physiology & Behavior 77, 677–682 (2002).

23 Cheney, P. D. & Fetz, E. E. Functional classes of primate corticomotorneuronal cells and their relation to active force. J. Neurophysiol. 44, 773–791 (1980).

24 Muir, R. B. & Lemon, R. N. Corticospinal neurons with a special role in precision grip. Brain Res. 261, 312–316 (1983).

25 Griffin, D. M., Hoffman, D. S. & Strick, P. L. Corticomotoneuronal cells are “functionally tuned”. Science 350, 667–670 (2015). 10.1126/science.aaa8035

26 Buchanan, T. S., Rovai, G. P. & Rymer, W. Z. Strategies for muscle activation during isometric torque generation at the human elbow. J. Neurophysiol. 62, 1201–1212 (1989).

27 Johanson, M. E., James, M. A. & Skinner, S. R. Forearm muscle activation during power grip and release. The Journal of hand surgery 23, 938–944 (1998).

28 Gribble, P. L. & Ostry, D. J. Compensation for interaction torques during single- and multijoint limb movement. J Neurophysiol 82, 2310–2326 (1999).

29 Charles, S. K. & Hogan, N. Stiffness, not inertial coupling, determines path curvature of wrist motions. J Neurophysiol 107, 1230–1240 (2012). 10.1152/jn.00428.2011

30 Formica, D. et al. The passive stiffness of the wrist and forearm. Journal of neurophysiology 108, 1158–1166 (2012). 10.1152/jn.01014.2011

31 Fromm, C. & Evarts, E. V. Pyramidal tract neurons in somatosensory cortex: central and peripheral inputs during voluntary movement. Brain Res 238, 186–191 (1982). 0006-8993(82)90781-8 [pii]

32 Westwick, D. T. & Kearney, R. E. Identification of physiological systems: a robust method for non-parametric impulse response estimation. Med Biol Eng Comput 35, 83–90 (1997).

33 Houk, J. C., Dessem, D. A., Miller, L. E. & Sybirska, E. H. Correlation and spectral analysis of relations between single unit discharge and muscle activities. J. Neurosci. Meth. 21, 201–224 (1987).

34 Sobinov, A. et al. Approximating complex musculoskeletal biomechanics using multidimensional autogenerating polynomials. PLOS Computational Biology 16, e1008350 (2020). 10.1371/journal.pcbi.1008350

35 Collinger, J. L. et al. High-performance neuroprosthetic control by an individual with tetraplegia. The Lancet 381, 557–564 (2013). 10.1016/S0140-6736(12)61816-9

36 Gilja, V. et al. A high-performance neural prosthesis enabled by control algorithm design. Nat Neurosci 15, 1752–1757 (2012). http://www.nature.com/neuro/journal/v15/n12/abs/nn.3265.html#supplementary-information

37 Agudelo-Toro, A., Michaels, J. A.Sheng, W.-A. & Scherberger, H. Accurate neural control of a hand prosthesis by posture-related activity in the primate grasping circuit. Neuron 112, 4115-4129. e4118 (2024).

38 Miller, L. E. et al. Correlation of primate red nucleus discharge with muscle activity during free-form arm movements. J. Physiol. London 469, 213–243 (1993).

39 Gibson, A. R., Houk, J. C. & Kohlerman, N. J. Magnocellular red nucleus activity during different types of limb movement in the macaque monkey. J. Physiol. 358, 527–549 (1985).

40 Davidson, A. G. & Buford, J. A. Motor Outputs From the Primate Reticular Formation to Shoulder Muscles as Revealed by Stimulus-Triggered Averaging. Journal of neurophysiology 92, 83–95 (2004). 10.1152/jn.00083.2003

41 Zaaimi, B., Dean, L. R. & Baker, S. N. Different contributions of primary motor cortex, reticular formation, and spinal cord to fractionated muscle activation. Journal of neurophysiology 119, 235–250 (2018). 10.1152/jn.00672.2017

42 Cohen, B. & Komatsuzaki, A. Eye movements induced by stimulation of the pontine reticular formation: evidence for integration in oculomotor pathways. Experimental neurology 36, 101–117 (1972).

43 Cannon, S. C. & Robinson, D. A. Loss of the neural integrator of the oculomotor system from brain stem lesions in monkey. Journal of neurophysiology 57, 1383–1409 (1987). 10.1152/jn.1987.57.5.1383

44 Delp, S. L. et al. OpenSim: open-source software to create and analyze dynamic simulations of movement. Biomedical Engineering, IEEE Transactions on 54, 1940–1950 (2007).

45 Truccolo, W., Eden, U. T., Fellows, M. R., Donoghue, J. P. & Brown, E. N. A point process framework for relating neural spiking activity to spiking history, neural ensemble, and extrinsic covariate effects. J Neurophysiol 93, 1074–1089 (2005).

46 Heinzl, H. & Mittlböck, M. Pseudo R-squared measures for Poisson regression models with over-or underdispersion. Computational statistics & data analysis 44, 253–271 (2003).

47 McFadden, D. Quantitative methods for analyzing travel behavior of individuals: some recent developments. (Institute of Transportation Studies, University of California Berkeley, 1977).

